# *Annulohypoxylon* sp. strain *MUS1*, an Endophyte isolated from *Taxus wallichiana* Zucc. produces Taxol and Other Bioactive Metabolites

**DOI:** 10.1101/2020.04.05.025858

**Authors:** Dhurva Prasad Gauchan, Heriberto Vélëz, Ashesh Acharya, Johnny R. Östman, Karl Lundén, Malin Elfstrand, M Rosario García-Gil

## Abstract

Endophytes are microbial colonizers that reside in plants by symbiotic association produces several biological classes of natural products. The current study focuses on the isolation and characterization of bioactive compounds produced by endophytic fungi isolated from the Himalayan yew (*Taxus wallichiana*) collected from the Mustang district of Nepal. The plant materials were collected from the Lower-Mustang region in the month of October 2016 and the endophytic fungi were isolated on artificial media from inner tissues of bark and needles. Antimicrobial and antioxidant activity, along with total phenolic- and flavonoid-content assays, were used in the evaluation of bioactivity of the fermented crude extracts along with the *in vitro* ability of the endophytes to produce the anticancer compound Taxol was analyzed. A total of 16 fungal morphotypes were obtained from asymptomatic inner tissues of the bark and needles of *T. wallichiana*. Among the 16 isolates, the ethyl acetate (EA) fraction of isolate *MUS1*, showed antibacterial and antifungal activity against all test-pathogens used, with significant inhibition against *Pseudomonas aeruginosa* ATCC 27853 (MIC: 250 µg/ml) and the pathogenic yeast, *Candida albicans* (MIC: 125 µg/ml). Antioxidant activity was also evaluated using 2,2-diphenyl-1-picrylhydrazyl (DPPH). At a concentration of 100 µg/ml, the % radical scavenging activity was 83.15±0.40, 81.62±0.11, and 62.36±0.29, for ascorbic acid, butylated hydroxytoluene (BHT) and the EA fraction of MUS1, respectively. The DPPH-IC_50_ value for the EA fraction was 81.52 µg/ml, compared to BHT (62.87 µg/ml) and ascorbic acid (56.15 µg/ml). The total phenolic and flavonoid content in the EA fraction were 16.90±0.075 µg gallic acid equivalent (GAE) and 11.59±0.148 µg rutin equivalent (RE), per mg of dry crude extract, respectively. Isolate *MUS1*, identified as an *Annulohypoxylon* sp. by ITS sequencing, also produced Taxol (282.05 µg/L) as shown by TLC and HPLC analysis. Having the ability to produce antimicrobial and antioxidant compounds, as well as the anticancer compound Taxol, makes *Annulohypoxylon* sp. strain *MUS1*, a promising candidate for further study given that naturally occurring bioactive compounds are of great interest to the pharmacological, food and cosmetic industries.

## Introduction

*Taxus wallichiana*, also known as Himalayan yew (Vernacular name: *Lauth-salla)*, is a medicinal plant found in hilly regions of Nepal. *T. wallichiana* grows on steep mountain slopes of altitude 1500-3000 meters, and mostly in areas not disturbed by man-made practices [1]. *Taxus* species are the source of a highly functionalized diterpene molecule called Taxol, first isolated from the stem bark of Western yew, *Taxus brevifolia* [2], and used in cancer treatment. This has led to the overharvesting of needles from the Yew trees and consequently, *Taxus* forests in the Himalayan regions are on the verge of extinction and now in the IUCN Red List. This has compelled an immediate search for alternative sources of Taxol in order to save these trees and maintain their biodiversity for future generations.

Endophytes are a group of microbial colonizers that reside in the inner tissue of plants [3]. These microorganisms form a symbiotic association with the host that can range from commensalistic to slightly pathogenic [4]. Some endophytes also have the ability to produce the same or similar secondary metabolites as their plant-host [5]. For example, the discovery that some endophytic fungi could produce Taxol opened the possibilities of using them as an alternative source to the *Taxus* plant [5, 6]. Since the initial report, more than 200 endophytic fungi isolated from plants have been reported to produce Taxol in culture media, albeit in small quantities or requiring optimization in some cases [7, 8].

Different biological classes of natural products, including antibacterial, antifungal, anticancer, antiviral as well as plant protective agents, have been reported to be produced by endophytes [4, 9]. However, the endophytic microbiota in plants still represents a largely untapped resource of natural products [10]. Therefore, the current study focuses on the isolation and characterization of bioactive compounds produced by endophytic fungi isolated from the Himalayan yew (*T. wallichiana*) collected from the Mustang district of Nepal.

## Materials and Methods

### Location of Study Area and Sample Collection

The *Taxus* plant materials were collected from *Taglung* forest of the Lower-Mustang region of Mustang district (Fig. S1, suppl. file), at similar altitudes (2722 m. to 2729 m.) and geographic locations (latitude 28.39022 to 28.39047 °N and longitude 83.37135 to 83.37240 °E) in the month of October 2016. The plant samples (i.e., healthy and asymptomatic bark and needles) from *T. wallichiana*, were packed in polythene bags and brought to the laboratory in an icebox. The samples were processed within 48 hours of collection and stored at 4 °C until isolation procedures were finished. The specimen of each sample was deposited at the Department of Biotechnology, Kathmandu University in Dhulikhel, Kavre, Nepal.

### Media and Microorganisms used

Potato dextrose agar (PDA, HiMedia) or Potato dextrose broth (PDB, HiMedia) was used for fungal isolation and routine growth of fungal endophytes, for genomic DNA (gDNA) extraction and secondary metabolite production. Gram-positive bacteria (*Streptococcus faecalis* ATCC 19433, *Staphylococcus aureus* ATCC 12600, and *Bacillus subtilis* ATCC 6633), Gram-negative bacteria (*Escherichia coli* ATCC 25922, *Salmonella enterica* ATCC 13076, *and Pseudomonas aeruginosa* ATCC 27853) and the yeast, *Candida albicans* (isolated from oral thrush of HIV patient and verified as *C. albicans*), were used as test-pathogens. The bacterial strains were obtained directly from the American Type Culture Collection (ATCC) and, *Candida albicans* was obtained from the Department of Microbiology, Dhulikhel Hospital of Kathmandu University, Dhulikhel, Nepal. All the pathogens were grown and maintained in Muller-Hinton Agar (MHA) or Muller-Hinton broth (MHB) [11].

### Sample Processing and Isolation of Endophytic Fungi

The isolation of fungi from inner tissues of needles and bark of *T. wallichiana* was carried out in two methods as previously described [12]. In the first one, the bark and needle samples were washed thoroughly with running tap water in order to remove the dust and external debris. The bark was cut into 1 cm x 1 cm x 0.5 cm pieces, while the needles were separated into groups of 4-6 needles and washed with sterile water before surface sterilization. Surface sterilization was carried out with 70% (v/v) ethyl alcohol for 2 minutes, followed by 0.2% mercuric chloride for 1 minute. The sterilized pieces of bark and needles were washed with sterile distilled water three times using separate containers. After washing with sterile water, bark and needles were dried using sterile blotting paper. A single needle was excised and was cut into 0.5 cm long pieces for each sample. The cut needles were placed over the PDA media (pH 5.6) horizontally --freshly-exposed tissue facing down-- or vertically, slightly dipping the breadth side of the needle into the media. The outer bark was removed using a sterile blade and inner sterile tissue was excised. The inner bark-tissue was cut into 0.5 cm x 0.5 cm pieces for each sample and placed over PDA media (pH 5.6) in a Petri-plate. The plates were sealed with parafilm and incubated at 25±2 °C for at least two weeks.

In the second method and as above, 0.5 cm × 0.5 × 0.5 cm sterile tissues were excised by removing the outer bark using a sharp blade. One to two bark pieces (0.5 cm × 0.5 cm) of each sample were ground into a paste with 2 ml of sterile-distilled water using a mortar and pestle. Similarly, two to three surface-sterilized leaf-needles were excised, blotted dry and ground. The maceration of bark and needle tissues was carried out in two steps. First, one ml of sterile distilled water was added to the sample and grounded with gentle force. In the second step, an additional one ml of sterile-distilled water was added and the sample was grounded well to make a paste/solution. About one ml of the grounded paste/solution was transferred to a Petri-plate and approximately 20 ml of molten PDA media was poured into the plate. The mixture was homogenized by gently moving the plate in the shape of the number eight. Once solid, the plates were sealed with parafilm and incubated at 25±2 °C for at least two weeks. No antibiotics were added to the media. Every single sample was processed by both methods. Hence, the processing of a single sample comprised four plates, two for intact tissue (one bark and one needle) and two for ground tissue (one for bark and one for needle). Fungal isolates obtained from both methods were checked for contamination. Finally, the pure fungal isolates were obtained by transferring the tissue-extruded hyphae filaments to another PDA media plate (pH 5.6) by using the hyphal tip method [1].

### Characterization of endophytic fungi

Morphotypes were assigned based on the phenotype of the fungal colony (e.g., color, shape, and size), mycelial mat characteristics in the growth media, as well as the presence of sexual or asexual structures seen under the microscope [13, 14]. Tape-lift mounts stained with Lactophenol cotton blue were visualized under the microscope [15].

Genomic DNA (gDNA) from the isolated fungal endophytes was extracted for molecular characterization using the Quick-DNA™ Fungal/Bacterial Miniprep Kit (Zymo Research Company) with little modification. For this, agar-plugs of isolated fungi grown in PDA were used to seed Erlenmeyer flasks containing 50 ml of PDB and incubated at 25±2 °C, shaking at 150 rounds-per-minute (rpm). After 3 days, mycelia were harvested by filtration and washed thoroughly with distilled water to remove any media and extracellular metabolites. The harvested mycelia (≈300 mg) was pulverized with liquid nitrogen using mortar and pestle and transferred to 2 ml Eppendorf tubes. Lysis Solution, 750 µl, was added to the tube containing the pulverized fungal mycelium powder and the samples were incubated at 65 °C for 1 hour. The remaining steps were followed as prescribed in the instruction manual provided with the kit.

The gDNA obtained was used for PCR to amplify the internal transcribed spacer (ITS) using primers ITS1 (F-5’-TCCGTAGGTGAACCTGCGG-3’) and ITS4 (R-5’-TCCTCCGCTTATTGATATGC-3’) to identify the endophytic fungi at the molecular level as previously described [16]. The 25 µl PCR-mixture consisted of 13.5 µL of PCR grade water, 1.25 µL of ITS1-F (10 µM), 1.25 µL of ITS4-R (10 µM), 2.5 µL of amplification buffer (10X), 0.75 µL of MgCl_2_ (50 mM), 0.5 µL of dNTP mix (10 mM), 0.25 µL of Taq DNA polymerase (5U/µL) and 5 µL of Template DNA. PCR was carried out in a thermocycler (BIO-RAD T100) under the following amplification conditions: an initial denaturing step at 94 °C for 5 min, followed by 35 amplification cycles of 94 °C for 1 min, 55 °C for the 40s, and 72 °C for 1 min and a final extension step at 72 °C for 10 min. The PCR products were visualized on 1% (w/v) agarose gel and aliquots were sent to Macrogen-Europe for sequencing. The obtained sequences were compared to existing DNA sequences in NCBI GenBank using BLASTn (megablast). The phylogenetic relations of the fungal endophytes were analyzed by MEGAX software [17].

### Production, Extraction, and Analysis of Secondary Metabolites

#### Production and Extraction

To test the production of intracellular and secreted extracellular secondary metabolites, a small-scale fermentation process was carried out as previously described [18]. Briefly, fungi were grown in 250 ml Erlenmeyer flasks containing 100 ml of PDB at 25 ±2 °C and shaken at 150 rpm. After 10 days, mycelia and liquid culture media were separated by filtration. The mycelia were homogenized with mortar and pestle and extracted with 10 ml each of either Methanol (MH), Ethyl acetate (EA), or Chloroform (CH). Similarly, the liquid filtrate was extracted with equal volumes (100 mL) of either EA, CH, or Hexane (HX), using a separatory funnel. The solvent-extracts obtained from both mycelia (10 ml) and culture-filtrates (100 ml) were filtered using Whatman filter paper No.1 and evaporated to dryness at room temperature under an exhaust fan. The dried-fractions extracted with each solvent (i.e., EA, MH, CH, or HX) were weighed and stored at 4 °C.

#### TLC analysis

Thin-layer Chromatography (TLC) analysis of the crude extracts was done according to Garyali, Kumar, and Reddy 2013, with little modifications [19]. The crude fractions were dissolved in EA (10 mg/ml) and 10 µl from each were spotted onto 20 × 20 cm TLC plates (Silica gel 60 F254, Merck) with at least one cm between the spots and placed in a binary mobile phase consisting of Chloroform: Methanol (9:1 v/v). Once the mobile phase reached the top, the TLC plate was removed, the solvent evaporated, and the plate was observed under short and long UV. Furthermore, the TLC plate was sprayed with 2% AlCl_3_ and again observed under short and long UV. A second TLC plate made in the same manner, and was sprayed with 0.04 mg/ml DPPH solution, incubated in the dark for 10 min, and observed under visible light.

#### Agar-well Diffusion Assay

Altogether, six human pathogenic-bacteria and one human yeast-pathogen were used in an agar-well diffusion (AWD) assay, to determine the antimicrobial activity of the crude extracts obtained from the fungal endophytes [11, 18, 20]. Stocks (10 mg/ml) for both the intracellular and extracellular-extracts were made using DMSO as a solvent. Antibiotic discs of gentamicin (10 µg/disc) and fluconazole (25 µg/disc), were used as positive controls against bacteria and yeast, respectively. Gram-positive bacteria (*Streptococcus, Staphylococcus, and Bacillus*), Gram-negative bacteria (*Escherichia, Salmonella, and Pseudomonas*) and the ascomycete yeast (*Candida albicans*) were grown overnight in MHB and adjusted to a McFarland turbidity standard of 0.5 (1 × 10^8^ Colony Forming Units (CFU)/ml). The test-pathogens were spread on MHA plates using an L-shaped spreader and allowed to dry for few minutes. Once dried, for each treatment, agar-wells were made with a sterile borer (6 mm inner diameter) and 50 μl of extract (10 mg/ml) were loaded into the agar-wells. Standard antibiotic disc of gentamicin or fluconazole (6mm in diameter) was placed on top of the media. DMSO-only (50 µl) was also included as a negative control. The diameter (mm) of the zone of inhibition (ZOI) was measured for each treatment and compared to the controls. The assay was carried out in triplicate for each test-pathogen.

#### MIC of selected fungal Crude Extracts

The minimum inhibitory concentration (MIC) was calculated using a macro-broth dilution method with little modifications [11]. *P. aeruginosa* and *C. albicans* were chosen as test-pathogens based on the greater ZOI values obtained in the AWD assay. The test-pathogens (0.5 McFarland turbidity standard, 1 × 10^8^ CFU/ml) were diluted with sterile-distilled water in the ratio of 1:100 to a concentration of approx. 10^6^ CFU/ml. Gentamicin sulfate (HiMedia), a broad-spectrum bactericidal agent, was used as a standard antibiotic for MIC assay. The antibiotic stock was filtered-sterilized (0.45 µm filter) and stored at 4°C until the use.

#### TLC-Bioautography assay

The antimicrobial activity of the extracts was also evaluated using the TLC-bioautography agar overlay method, which is a combination of contact and direct bioautography [21]. After the TLC plate was prepared as described above, the plate was placed in a 90 mm Petri-plate and covered with molten MHA medium to a thickness of 2-3 mm. Once solid, the agar was overlaid with 10^6^ CFU/ml of *P. aeruginosa*. The plates were incubated at 37±2 °C for 18-24 hours. After incubation, the ZOI, which can be seen as clear spots against a purple background, was visualized by staining with 3-(4,5-dimethylthiazol-2-yl)-2,5-diphenyl tetrazolium bromide (MTT) solution (0.5 mg ml^−1^).

#### Radical scavenging assay

A DPPH-radical scavenging assay using 2,2-diphenyl-1-picrylhydrazyl (DPPH; Sigma-Aldrich) was carried out as previously described [22] with some modifications. The DPPH reagent was freshly prepared in Methanol (0.04 mg/ml). Ascorbic acid (used as a positive control), and the EA fraction were prepared in methanol in the concentration range of 5, 10, 20, 40, 60, 80, 100 and 120 µg/ml. One ml of the standards, or the EA fraction, was mixed with three ml of the DPPH solution (1:3) in a glass test-tube. After mixing, the test tubes were incubated in the dark at 27°C for 10 min. The absorbance at 517 nm was recorded using a UV-Vis Spectrophotometer (Shimadzu, UV-1800) and Methanol was used to blank the instrument. The assay was repeated three times. The percent scavenging ability for DPPH-radicals was calculated as described in Pan et al. 2017 [22]. Additionally, a measure of half-maximal inhibitory concentration (IC_50_) for 50% DPPH scavenging activity was carried out for standards and EA fraction of MUS1 by fitting a linear regression to DPPH scavenging curves respectively.

#### TPC and TFC Assays

The total phenolic content (TPC) and total flavonoid content (TFC) of the fungal crude extracts obtained using the solvents EA, CH, and HX, were determined as described [22]. However, the assay volumes were halved (total volume = 12.5 ml) and methanol was used to dissolve the standards i.e., gallic acid (Himedia), rutin (Himedia), and the crude extracts (1 mg/ml). For the TPC assay, the absorbance was measured at 733 nm, while the TFC assay was measured at 507 nm using a UV-Vis Spectrophotometer (Shimadzu, UV-1800). The TPC from the extracts was determined based on the calibration curve obtained from the gallic acid standards (i.e., 1, 2, 4, 6, 8, 10, 12, and 16 µg/ml), and expressed as µg of gallic acid equivalents (GAEs) per mg of dry fungal extract. Similarly, the calibration curve from the rutin standards (i.e., 5, 10, 20, 40, 60, 80, 100, and 120 µg/ml), was used to calculate the TFC and expressed as µg of rutin equivalents (REs) per mg of dry fungal extract.

### Taxol Analysis

#### Preparative TLC

TLC was used to test for the presence of Taxol in the EA fraction obtained from crude extracts of the endophytic fungal isolates [23]. Twenty µl of the EA fraction (10 mg/ml) were spotted onto 20 × 20 cm TLC plates (Silica gel 60 F254, Merck) with at least one cm between the spots and placed in a quaternary mobile phase consisting of Benzene: Chloroform: Acetone: Methanol (20:92.5:15:7.5). Once the mobile phase reached the top, the TLC plate was removed. The standard anticancer compound, Taxol (Paclitaxel; Sigma-Aldrich) was also included and used for comparison. The bands horizontally in-line with the Taxol standard were visualized under short UV and marked with a pencil. The pencil-marked spots were scraped off the plates using a sharp blade and eluted using acetone-methanol (1:1) mixture followed by centrifugation at 10,000 rpm for 10 minutes. After centrifugation, the supernatant was taken and evaporated to dryness under reduced pressure and dissolved in methanol for the HPLC analysis of Taxol.

#### RP-HPLC Analysis

Reverse-phase HPLC (RP-HPLC) analysis and quantification of Taxol was done using a reverse-phase C18-column (Shim-pack GIST C18, 5 µm, 150 × 4.6 mm I.D.) in a Shimadzu LC-2030 instrument with UV absorbance at 227 nm as described elsewhere [24, 25] with some modifications. For this, 15 µl of the sample was injected into the column at a column temperature of 30 °C. The separation was carried out at a flow rate of 1 ml min^-1^ with a gradient acetonitrile-water (v/v) system (50% of acetonitrile for 0-10 mins, 90% of acetonitrile for 10-12 mins, again 50% of acetonitrile for 12-15 mins and stop after 15 mins). A standard curve (R^2^ = 0.99) was generated using authentic Taxol (Paclitaxel; Sigma-Aldrich) solutions (i.e., 5, 10, 15, 25, 50, 100 µg/ml) and used to calculate the amount of Taxol present in fermentation cultures as described in Liu et al. 2009 [25].

#### Statistical Analysis

Statistical analysis was carried out using GraphPad Prism version 8.0.0 for Windows, GraphPad Software, San Diego, USA, for t-test, ANOVA and Post Hoc tests. The t-test and one-way ANOVA followed by Tukey’s Post Hoc test with p < 0.05 were used to analyze the significant differences between the results obtained.

## Results

### Endophytic Fungi from Himalayan Yew

A total of 16 fungal morphotypes were obtained from asymptomatic inner tissues of bark and needles of *T. wallichiana* using PDA media.

### Identification of isolate MUS1 from Himalayan Yew

Fungal genera identified by morphological characterization included *Alternaria, Aspergillus, Fusarium, Mucor, Penicillium*, and *Trichoderma*. Some of the endophytic isolates were unidentified. An endophytic fungus isolated among the 16 different morphotypes, initially named as isolate *MUS1*, exhibited bioactive-compound activity and was chosen for further study. The isolate showed a creamy-white flocculent mycelium in the dorsal-side of the PDA plate and yellowish-brownish-black taint on the ventral-side of the PDA plate after 10 days of incubation. No signs of sporulation or conidia were seen. *MUS1* was identified as *Annulohypoxylon* sp. (Fig. 2) and the ITS sequence has been deposited in GenBank under accession number MN699475.

**Figure 1:**
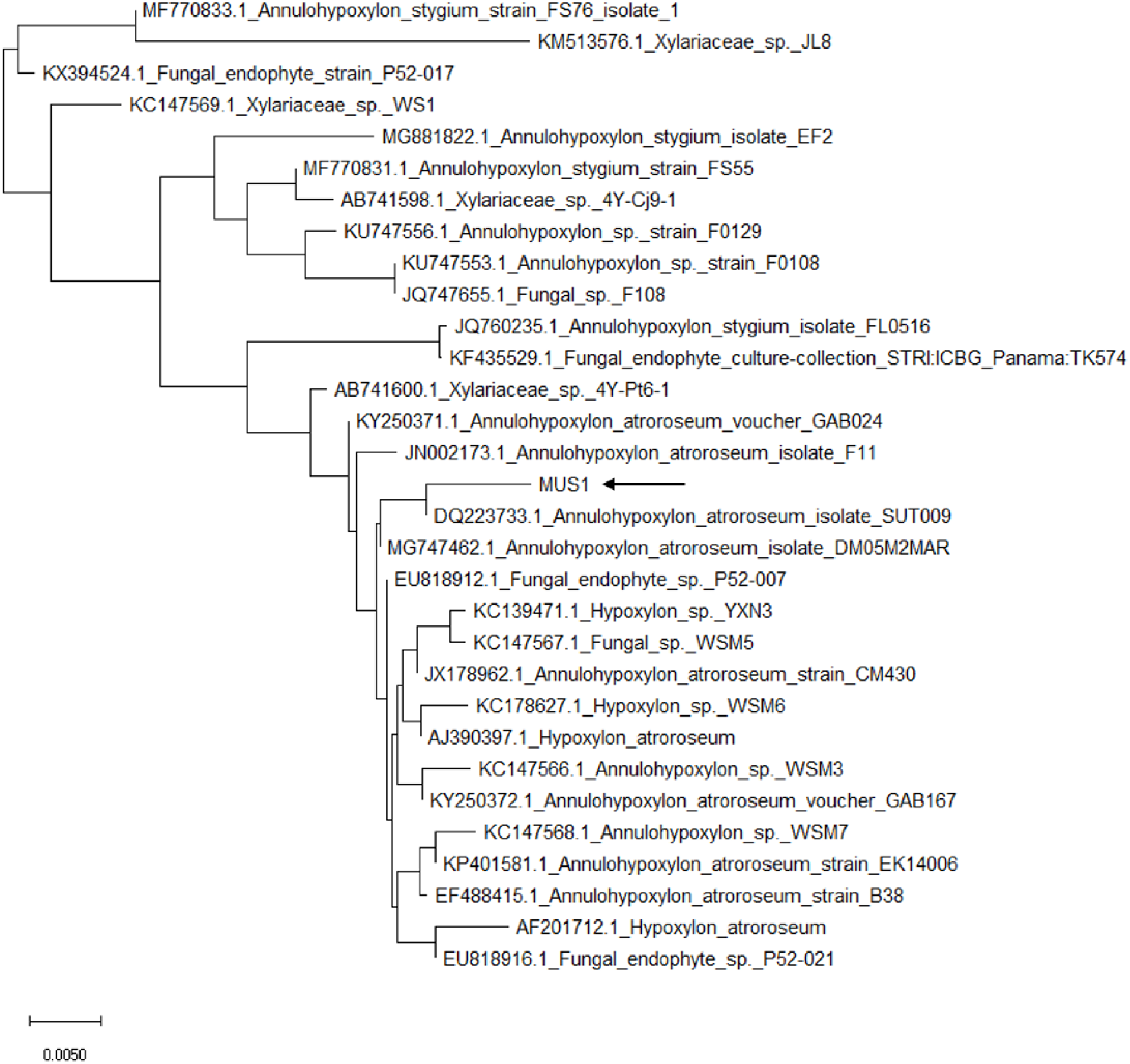
Phylogenetic analysis of ITS1-5.8S-ITS2 sequences from the endophytic fungi of Himalayan yew isolated from Mustang districts of Nepal using Neighbor-joining (NJ) method. The sequence of endophytic fungi under study (isolate *MUS1*) marked with symbol MUS1 (pointed with arrow) with bootstrapping of 1000 replicates (conducted in MEGAX software).

**Figure 2.**
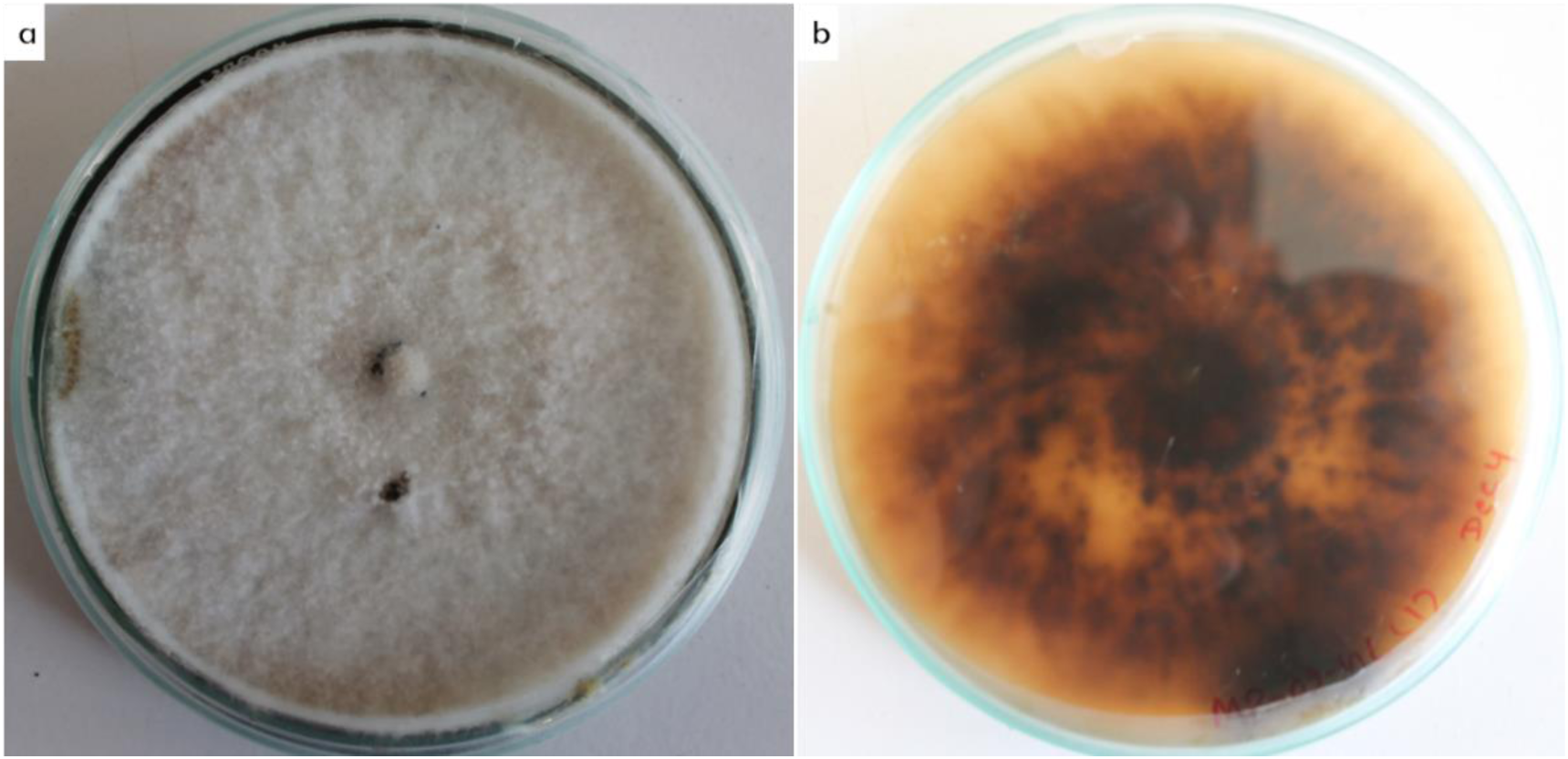
Two-week-old culture of fungal isolate, *MUS1*, later identified as *Annulohypoxylon* spp. Grown on Potato dextrose agar (PDA) media after 10 days of incubation, (a) dorsal side – a creamy-white flocculent mycelium, and (b) ventral side – yellowish-brownish-black color. No signs of sporulation or conidia were seen under the microscope.

### Antimicrobial activity

The EA fraction from the 16 fungal endophytes isolated, was analyzed for antimicrobial activities against seven human pathogens using the AWD assay. Only the extracellular EA-fraction of isolate *MUS1* showed antimicrobial activity against all the pathogens tested. Two isolates showed activity against the bacterial pathogens but had less activity as compared to *MUS1*. Other isolates were unable to produce any antimicrobial activity (results not shown). Out of the three solvents used in the extraction (i.e., HX, EA, or CH), the EA fraction (500 µg/well) had significant antimicrobial activity compared to the CH or HX fractions (Tukey’s post-hoc test, P < 0.05; results not shown), suggesting that the compound or compounds responsible for the antimicrobial activity were soluble in EA. However, no activity was seen for the intracellular fractions for any of the solvents used including EA in the extraction (results not shown). Interestingly, the yield of the extracellular metabolites was less in comparison to intracellular metabolites from isolate *MUS1* for all the solvents used (Table 1).

**Table 1:** Metabolite extraction from isolate *MUS1*.

| Solvent fraction | Intracellular Metabolites (mg)# | Extracellular Metabolites (mg)# |
| --- | --- | --- |
| EA | 29.5 [7.04] | 40.1 [0.0401] |
| CH | 17.2 [4.11] | 9.8 [0.0098] |
| HX | N/a | 8.6 [0.0086] |
| MT | 80.9 [19.31] | N/a |

There was a significant difference in the ZOI between the EA fraction and the DMSO (control) for all the seven pathogens tested (t-test, P < 0.05, Fig. 4). Furthermore, comparing the EA fraction with the antibiotic gentamicin (positive control), there was no significant difference in ZOI against *P. aeruginosa* (10 µg gentamicin vs. 500 µg crude extract). This would suggest that the compound present in the EA fraction was as good as the standard antibiotic gentamicin used against *Pseudomonas aeruginosa* (t-test, P > 0.05, Fig. 5). Moreover, the EA fraction showed a significant ZOI against the yeast pathogen, *C. albicans*, in comparison to fluconazole (positive control) or DMSO (negative control) (Figures 5 and 6, t-test, P < 0.05). These results showed that the extracellular crude extract from the fungal endophyte, isolate *MUS1*, contains antimicrobial compounds.

**Figure 3.**
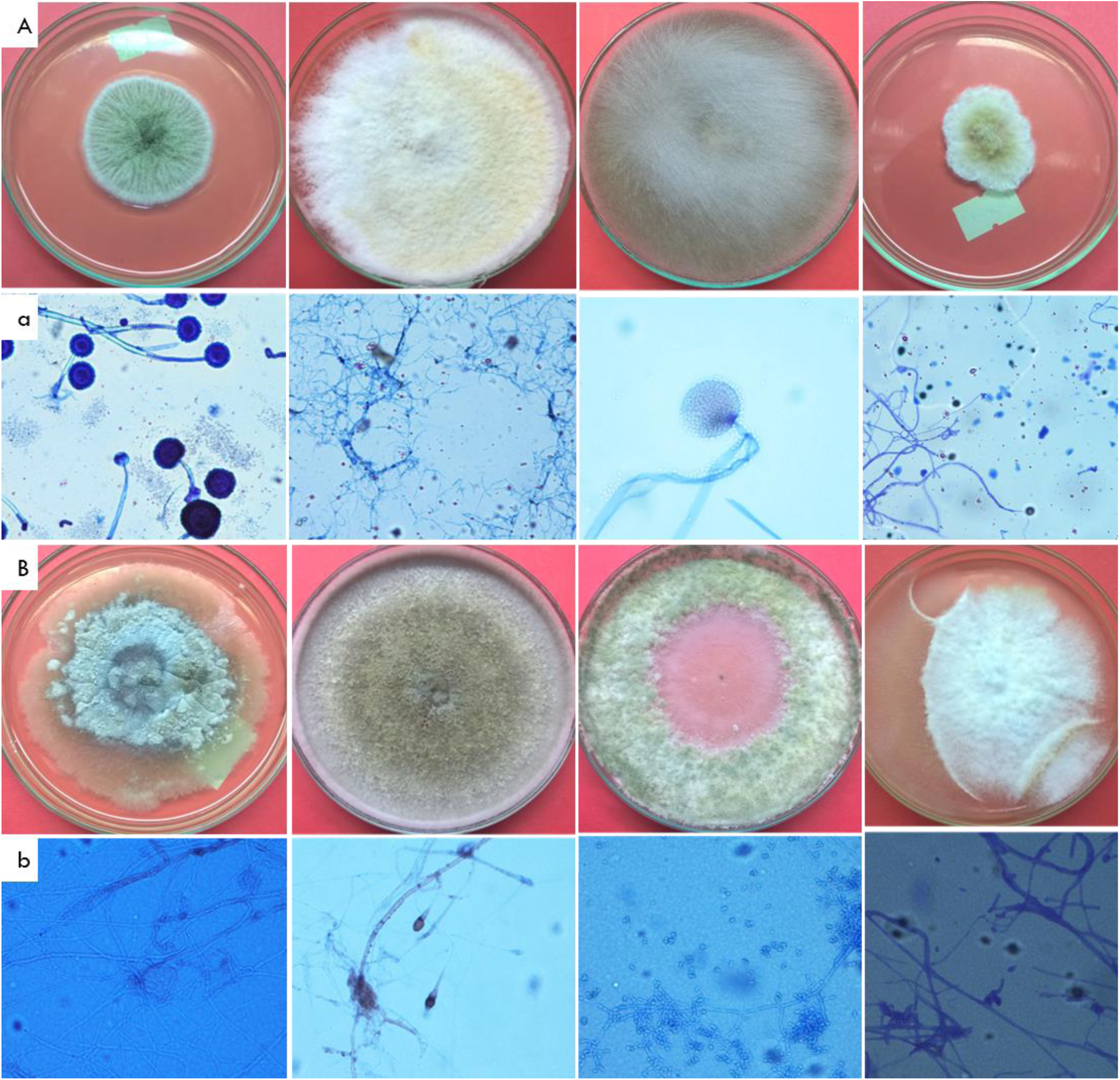
Morphological characteristics of two-week-old cultures of fungal endophytic isolates from *Taxus wallichiana*, (A, B) dorsal view of fungal isolates in PDA media, and (a, b) respective microscopic structures seen under the compound microscope (400X).

**Figure 4.**
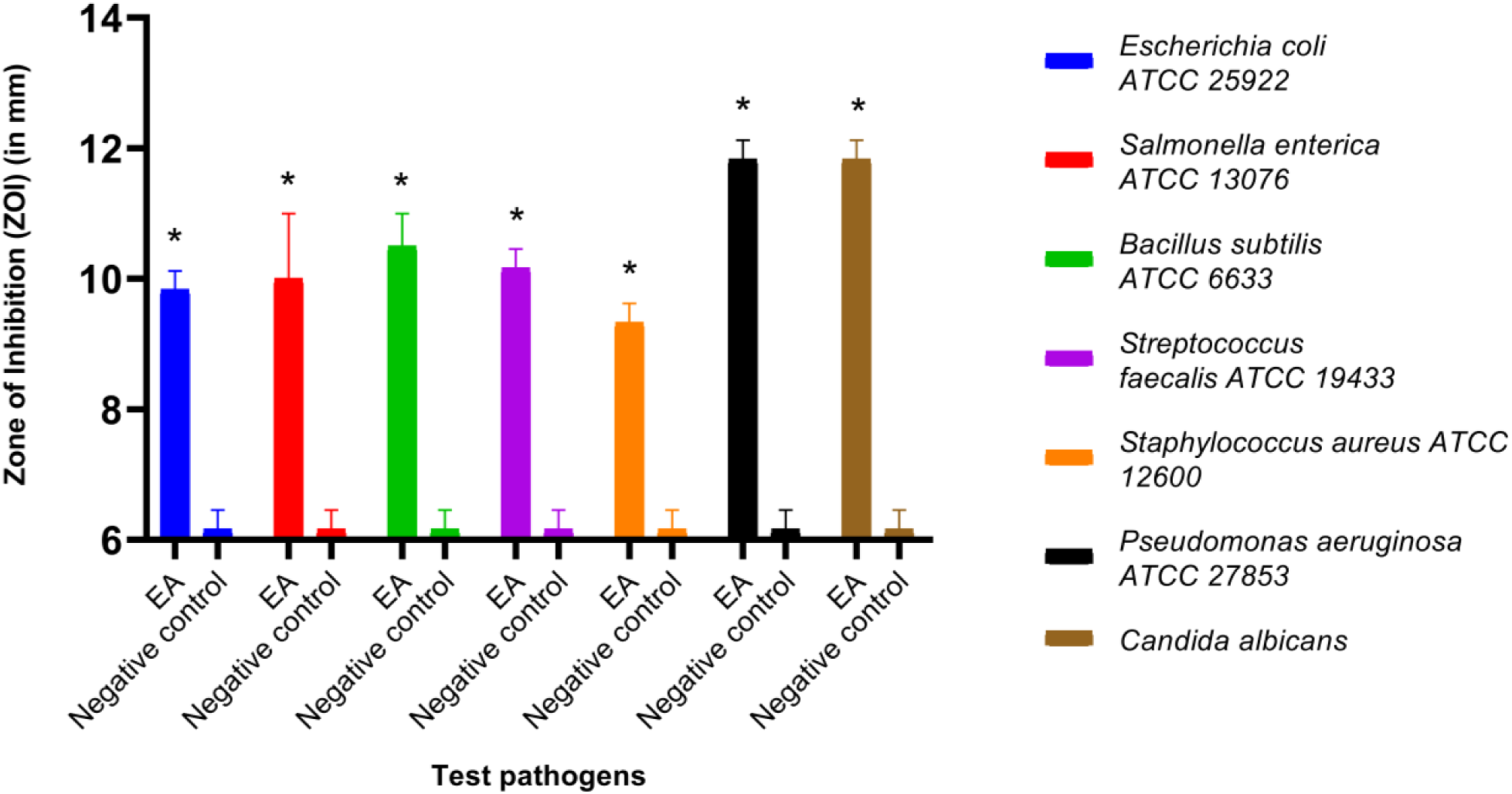
Zone of inhibition (ZOI) produced by the EA fraction (500 µg/well) from isolate *MUS1* and dissolved in Dimethyl sulfoxide (DMSO), compared to the DMSO alone. The EA fraction produced bigger ZOI for the seven human pathogens tested, compared to the DMSO. Data are presented as Mean ± SD (n=3). A ZOI value of 6 (size of an agar-well) was assigned for the diameter of the agar-well for when no inhibition was seen. *represents the significant difference in the ZOI between EA extract and Negative control on the test pathogens at P < 0.05, meaning effective control of pathogen in comparison to the negative control.

**Figure 5.**
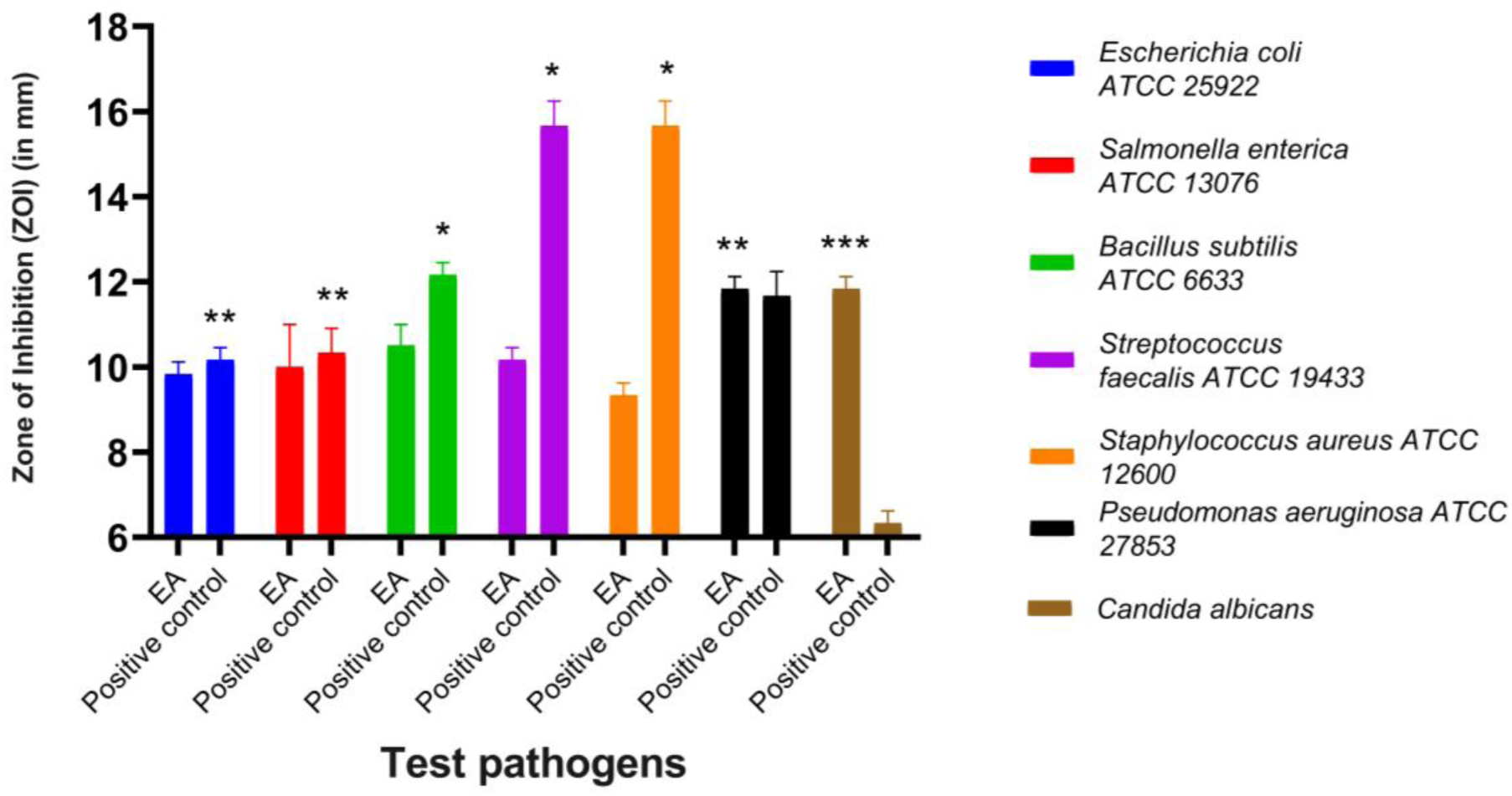
Zone of inhibition (ZOI) produced by the EA fraction (500 µg/well) isolated from isolate *MUS1*, compared to gentamicin and fluconazole for the seven human pathogens tested. Gentamicin (10 µg/disc) and fluconazole (25 µg/disc) were used as positive controls against bacteria and yeast, respectively. Except for *Streptococcus* and *Staphylococcus* spp., the EA fraction performed as well or better than the controls. Data are presented as Mean ± SD. A ZOI value of 6 (size of an agar-well) was assigned for the diameter of the agar-well when no inhibition was seen. *represents the significant difference in the ZOI between Positive control (Gentamycin disc, 10 µg) and EA extract (500 µg) on the bacterial pathogens at P < 0.05 (has antibacterial activity but less potent as compared to the antibiotic used as positive control). ** represents no significant difference in the ZOI between Positive control (Gentamycin disc, 10 µg) and EA extract (500 µg) on the bacterial pathogens at P > 0.05 (has antibacterial activity having similar potency to the antibiotic used as positive control). ***represents a significant difference in the ZOI between EA extract and Positive control (Fluconazole disc) on the yeast pathogen at P < 0.05 (has antifungal activity and more potent as compared to the antibiotic used as positive control).

**Figure 6.**
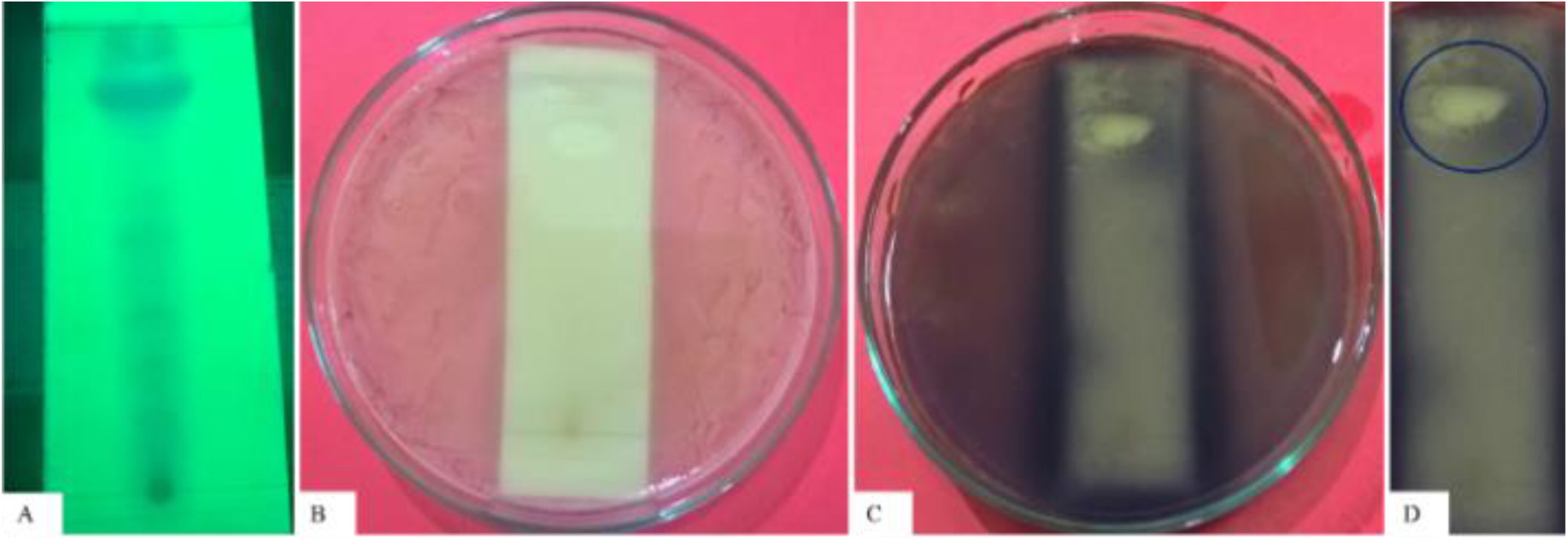
TLC-bioautography assay against *Pseudomonas aeruginosa* ATCC 27853 (**A**): TLC profile of metabolites under long UV light, (**B**): Bioautography assay, (**C**): Sprayed with MTT solution, and (**D**): Colorless inhibition zone (marked with circle), showing the active component in the crude extract responsible for antimicrobial activity.

The EA fraction that showed a potent antimicrobial activity during the AWD assay, was further analyzed for MIC against the human-pathogens, *P. aeruginosa* and *C. albicans*. The EA fraction of isolate *MUS1*, showed a MIC of 250 and 125 µg/ml against *P. aeruginosa* and *C. albicans*, respectively. The MIC for gentamicin against *P. aeruginosa* was 3.125 µg/ml. No standard MIC was analyzed for *C. albicans* as it was resistant to the antifungal agent, fluconazole.

The inhibitory activity of the EA fraction from isolate *MUS1*, as shown in the TLC bioautography assay, was attributed to one major constituent, spot 8 (R_f_ = 0.82), which gave a clear zone of inhibition seen against a purple background (Table 2 and Fig. 6). As expected, dead bacteria cannot reduce the MTT, producing the characteristic color change from yellowish to purple.

**Table 2:** TLC-bioautography assay against *Pseudomonas aeruginosa* ATCC 27853.

| Spot Number | R<sub>f</sub> value | 657<br>ZOI |
| --- | --- | --- |
| 1 | 0.25 |  |
| 2 | 0.34 |  |
| 3 | 0.4 |  |
| 4 | 0.45 |  |
| 5 | 0.51 |  |
| 6 | 0.55 |  |
| 7 | 0.6 |  |
| 8 | 0.82<sup>#</sup> | Positive |
|  | 0.98 |  |

<sup>#</sup> Inhibitory activity, after TLC separation of the EA fraction from isolate *MUSI*, was attributed to one major constituent, spot 8 (R<sub>f</sub> = 0.82), which gave a clear zone of inhibition (ZOI) seen against a purple background. R<sub>f</sub> = retardation factor.

### Metabolite analysis

TLC analysis of crude extracts from isolate *MUS1* extracted using various solvents showed a diversity of extracellular metabolites. However, the number of metabolites was higher for the EA fraction (R_f_ = 0.25 to 0.98) in comparison to the CH and HX fractions (Fig. 7). Furthermore, when the TLC plate was sprayed with a 2% AlCl_3_ solution, a bluish-green fluorescence spot could be seen under short-UV suggesting the presence of flavonoid compounds. The second TLC plate made in the same manner but sprayed with the DPPH solution also showed yellow spots on the fluorescent regions (Fig. 7). This suggests that extracellular metabolites from isolate *MUS1* are potentially rich in bioactive compounds including antioxidants.

**Figure 7.**
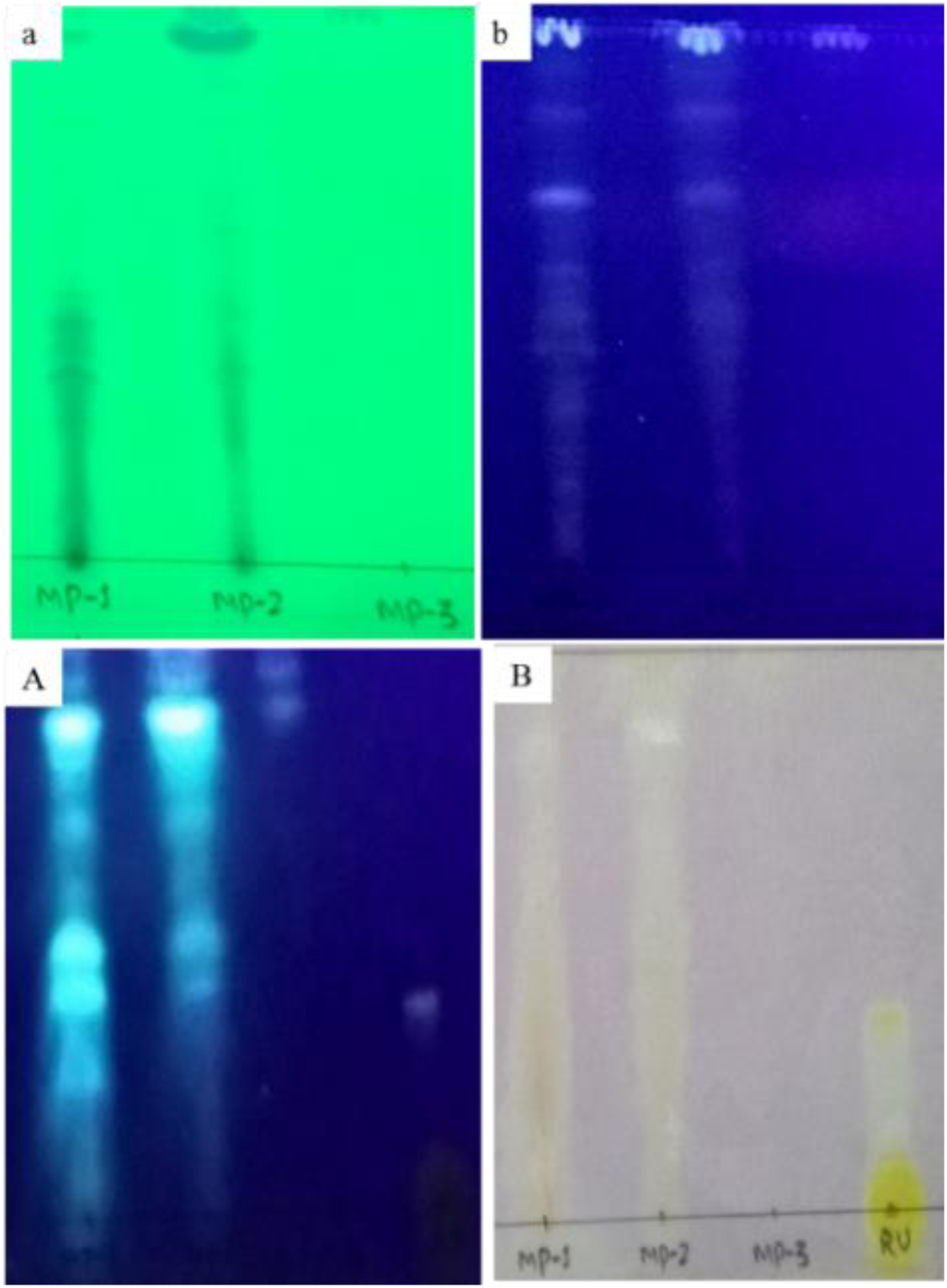
Thin-layer chromatography (TLC) plates of crude fractions from isolate *MUS1*. The three different fractions, **MP1** (extracted with ethyl acetate (EA)), **MP2 (**extracted with chloroform (CH)) and **MP3** (extracted with hexane (HX)), in each plate were run in a solvent mixture of chloroform: methanol (9:1) and visualized using different methods: (**a**) - under long UV light, (**b**) - under short UV light, (**A**)-TLC chromatogram after spraying with 2% AlCl_3_ under short UV light, and (**B**)-TLC chromatogram after spraying with 0.04 mg/ml DPPH under visible light. RU denotes the standard flavonoid compound, rutin. The chromatograms in plates **a** and **b** show that the number of metabolite bands is decreasing with decreasing polarity of the solvents used (i.e., higher for EA (**MP1**) in comparison to CH (**MP2**) and HX (**MP3**)). Similarly, fluorescence, after spraying 2% AlCl_3_, is higher in **MP1** than **MP2** and **MP3** (plate **A**).

### Antioxidant activity

The antioxidant activity of the EA fraction of isolate *MUS1* was evaluated by the DPPH-radical scavenging assay. As shown in the graph (Fig. 8), an increase in the concentration of the EA fraction, increased the percentage of DPPH-radical inhibition. For example, at 100 µg/ml concentration, the % DPPH-radical scavenging activity was 83.15±0.39, 81.62±0.06 and 62.36±0.29 for ascorbic acid, BHT and the EA fraction from isolate *MUS1*, respectively (Fig. 8). Similarly, the concentration required for 50% inhibition of DPPH free radical (IC_50_) was 56.15, 62.87 and 81.52 µg/ml for ascorbic acid, BHT and the EA fraction of isolate MUS1, respectively (Fig. 9).

**Figure 8.**
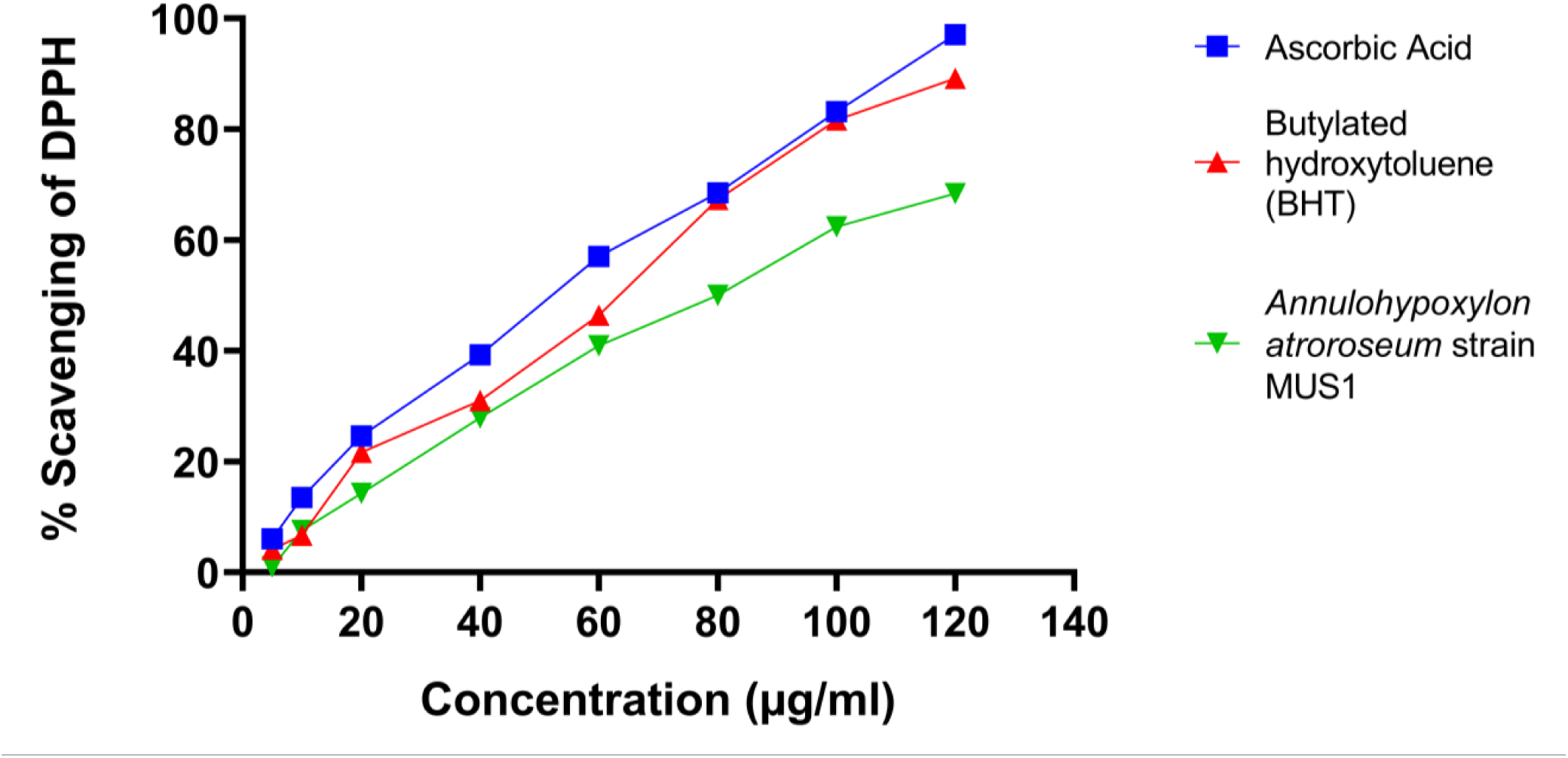
DPPH-radical scavenging assay. The % scavenging ability of the EA fraction from isolate *MUS1*, compared to the standard compounds, Ascorbic acid, and BHT, is shown. Data are presented as Mean ± SD.

**Figure 9:**
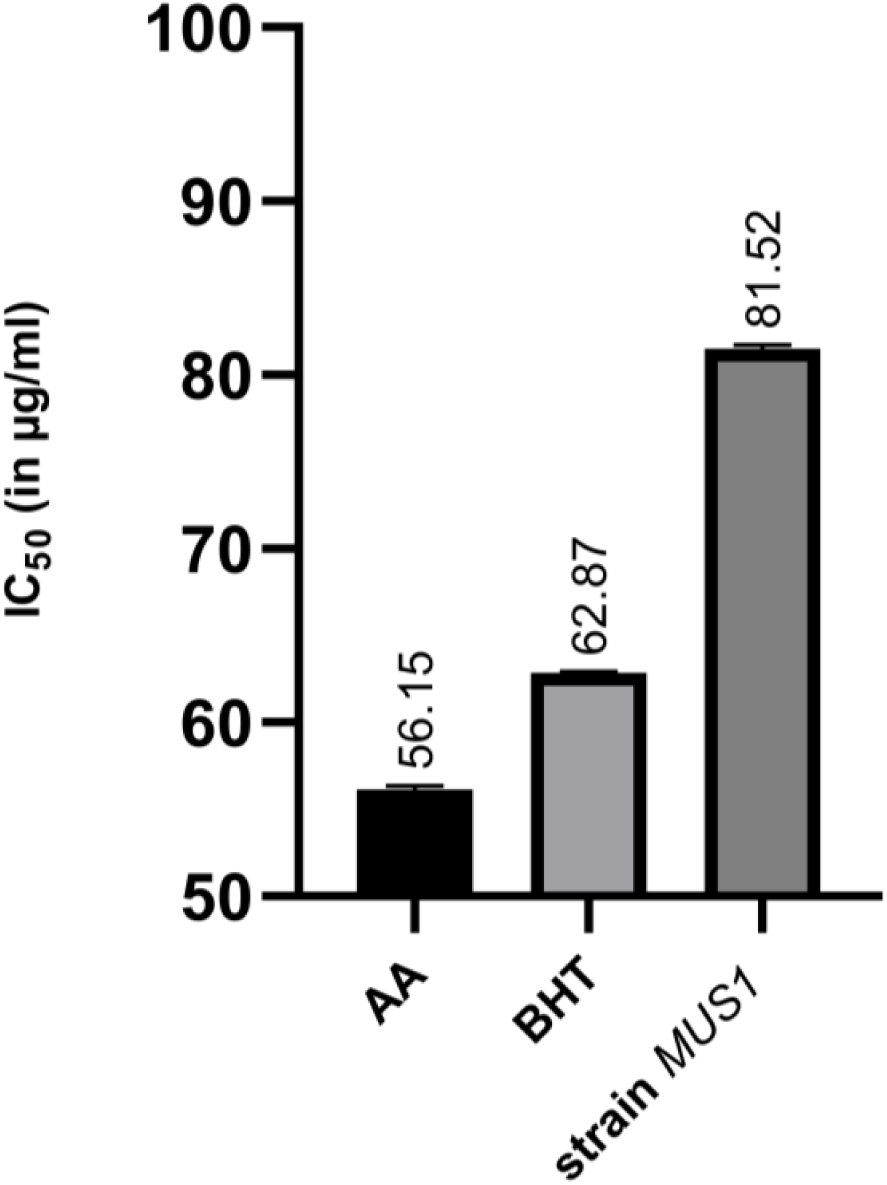
Graphical representation of the measure of half-maximal inhibitory concentration (IC_50_) for 50% DPPH scavenging activity (Mean ± SD) for Ascorbic acid (AA), Butylated hydroxytoluene (BHT) and isolate MUS1 under investigation.

### Phenol and Flavonoid content

The TPC and TFC were analyzed for extracts obtained from three different solvents and expressed in gallic acid equivalents (GAE) and rutin equivalents (RE) in µg per mg of dry fungal crude extract, respectively (Table 3). Both, the TPC and TFC, were higher in the EA extract compared to the CH and HX extracts. Moreover, taking into account the DPPH-radical scavenging activity of the EA extract (Fig. 8), it would suggest that the antioxidant activity of the EA extract could be due to phenolic and flavonoid compounds present in the extract.

**Table 3:** Total phenolic and flavonoid content for isolate *MUS1*.

|  | Total phenolics | Total flavonoids |
| --- | --- | --- |
| Solvent fraction | ($\mu\text{g GAE/mg dry extract}$) | ($\mu\text{g RE/mg dry extract}$) |
| Ethyl Acetate | 16.90±0.075 | 11.59±0.148 |
| Chloroform | 10.06±0.121 | 10.57±0.074 |
| Hexane | 0.93±0.068 | 1.97±0.074 |

Values are means of three biological replicates (n =3).

### Taxol analysis

TLC analysis of the EA fraction of isolate *MUS1*, showed two putative fungal-Taxol spots (R_f_ = 0.75 and 0.76) sharing the same R_f_ value as the Taxol standard (R_f_ = 0.75) (see Table 4). All three spots, which were scrapped and purified from the TLC plate, were further analyzed by RP-HPLC equipped with a detector system. Both, the authentic Taxol standard as well as the sample from isolate *MUS1*, had a retention time of 5.85 min (Fig. 10). The concentration of Taxol obtained was calculated to be 282.05 µg/L.

**Table 4.** TLC analysis of the ethyl acetate fraction for Taxol production by isolate *MUS1*.

| R<sub>f</sub> values |  |  |
| --- | --- | --- |
| Spot Number | *MUS1* | Taxol Standard |
| 1 | 0.18 |  |
| 2 | 0.31 |  |
| 3 | 0.75<sup>#</sup> | 0.75<sup>#</sup> |
| 4 | 0.76<sup>#</sup> |  |

**Figure 10:**
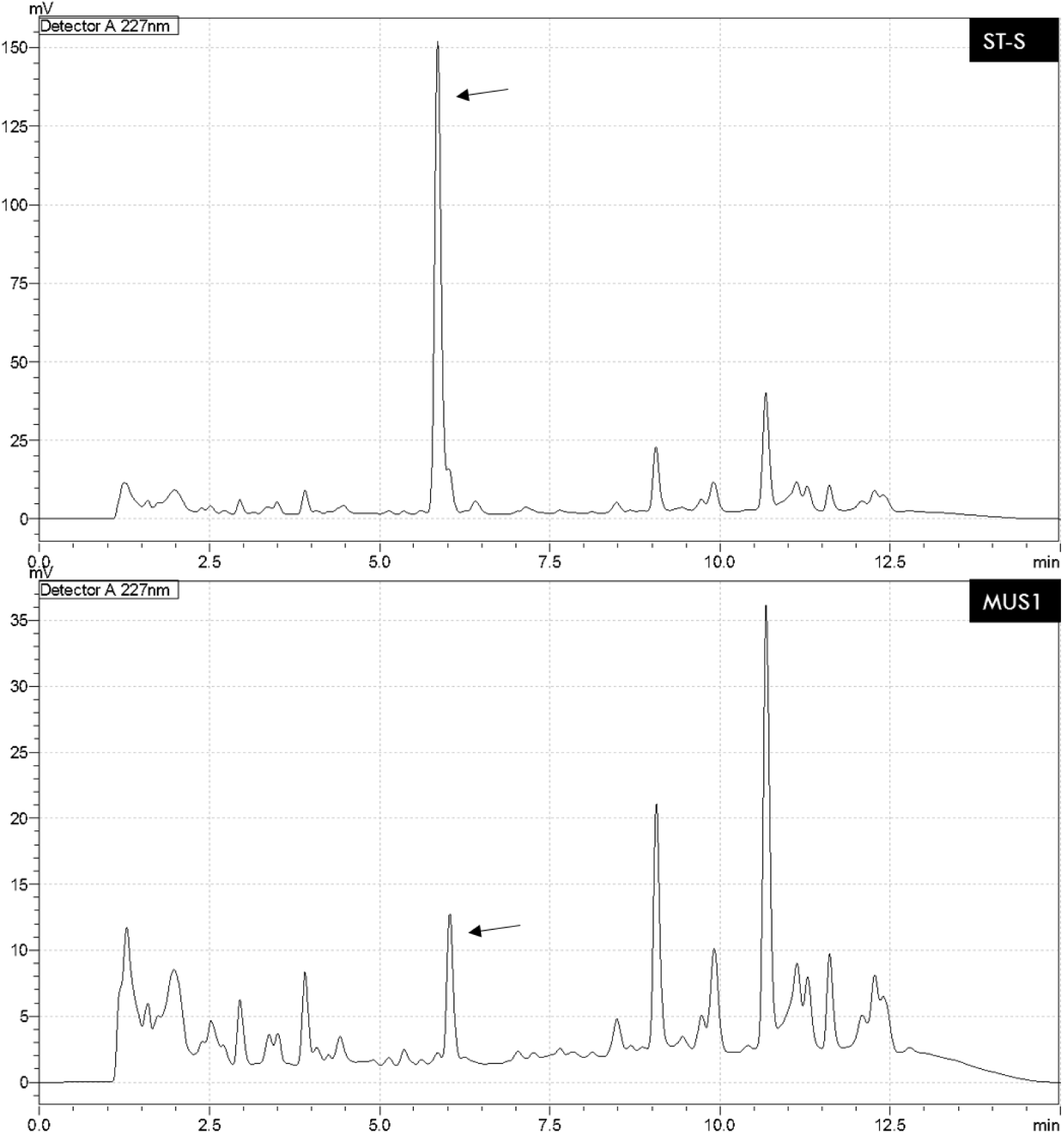
HPLC chromatograms showing the same retention time (5.85 min, shown with arrow) for the ST-S and the MUS1 spots obtained in the TLC assay. Both spots were scrapped from the silica, purified and analyzed using reverse-phase HPLC. **ST-S** is the authentic Taxol standard while **MUS1** corresponds to the putative Taxol compound obtained from the EA fraction of isolate *MUS1*.

## Discussion

Forest trees confer the largest niche for endophytic fungal diversity[26]. In fact, it is known that endophytic fungi isolated from plants show antimicrobial and antioxidant activities [18, 20, 22], including other bioactivities such as enzyme activity [27], volatile biocontrol agents [28] and volatile compounds having fuel potentials [9, 29, 30]. Moreover, endophytic fungi have the ability to produce similar or unique bioactive secondary metabolites compared to host plant metabolites [4, 5, 26, 31].

The Himalayan yew, *T. wallichiana*, a source of the anticancer drug Taxol and its related analogs [2], also harbors several fungal endophytes, some of which have been reported to produce Taxol in the artificial medium [8, 32]. Despite the relatively large number of studies that have been done on endophytic fungi isolated from *T. wallichiana*, the focus has been on Taxol and therefore, little information is available on antimicrobial and antioxidant properties of these fungal isolates. Recently, it was reported that crude extracts obtained with ethyl acetate from a fermented broth of *Fusarium* sp. isolated from *T. baccata* showed significant antimicrobial activity against all pathogens tested (i.e., *Bacillus subtilis, Staphylococcus epidermis, Staphylococcus aureus, Klebsiella pneumoniae, Shigella flexneri, Escherichia coli, Candida albicans*, and *Candida tropicalis*) [33]. Similarly, among the 16 endophytic fungi isolated during our study, *Annulohypoxylon* sp. strain *MUS1* showed a potential for the production of antimicrobial compounds. The AWD assay used in our study alluded to the presence of a compound or group of compounds in the EA fraction that had antimicrobial activity against several human pathogens, which included *Streptococcus faecalis, Staphylococcus aureus, Bacillus subtilis, Escherichia coli, Salmonella enterica, Pseudomonas aeruginosa*, and the yeast *Candida albicans*. Although there is a report for a *Hypoxylon* sp., isolated from *Persea indica*, producing volatile organic compounds that have potential as fuels or fuel additives [30], to our knowledge, this is the first report on the isolation and related bioactivities of an *Annulohypoxylon* sp. from *Taxus* sp.

Along with antimicrobial activity, fungal endophytes also show good promise as sources for compounds having antioxidant capacities. For example, it was recently reported that fungal endophytes isolated from the bulbs of *Fritillaria unibracteata* var. *wabuensis*, have a new potential as a natural resource for antioxidant compounds [22]. Likewise, the radical scavenging assay using DPPH showed that the antioxidant activity of the EA fraction from isolate *MUS1*, is comparable to standard antioxidants, *viz*., ascorbic acid and BHT. The EA fraction also showed the greatest amount for phenolics and flavonoids content (Table 3). Naturally occurring phenolic and flavonoid compounds are of great interest to the pharmacological industries as well as the food or cosmetic industries, as these compounds exhibit many biological functions including anti-inflammatory, antiallergic, hepatoprotective, antithrombotic, antiviral, anticarcinogenic and vasodilatory actions [34, 35].

Since the first report in 1993 for endophytic-Taxol production, many studies have been published showing that some endophytic fungi isolated from *Taxus* spp. are also capable of producing Taxol [8, 32]. Therefore, the 16 endophytic fungi isolated during our study were also evaluated for the production of Taxol using TLC. Only one fungus, isolate *MUS1*, showed the ability for the production of Taxol and this was confirmed using RP-HPLC analysis. The concentration of Taxol obtained was calculated to be 282.05 µg/L, which agrees with the published values for other fungi able to produce Taxol [8, 32].

Interestingly, our results during the initial screening with PCR primers used to amplify the genes *dbat* and *bapt* were negative (results not shown). These genes encode the enzymes 10-deacetylbaccatin III-10-O-acetyl-transferase and C-13 phenylpropanoid side-chain-CoA acyltransferase, respectively, that are involved in the Taxol biosynthetic pathway of *Taxus* spp. Although these PCR primers have been used to screen gDNA from fungi deemed able to synthesize the compound [36-38], the reverse PCR primer designed by Zhang and colleagues [39] for the *dbat* gene, is based on the plant sequence (GenBank No. EF028093) and is an intron-spanning primer. Multiple sequence alignment of the only four fungal-*dbat* sequences available in GeneBank (i.e., KP136287; EU883596; GU392264; EU375527, which also includes the one for *Cladosporium cladosporioides* MD2), and the plant sequence is almost 100% identical (data not shown). Therefore, cross-contamination between endophytic fungi and host DNA, as mentioned by Yang et al., 2014 [40], could account for the high similarity between these sequences. In the recent transcriptome analysis of the Taxol-producing fungus, *C. cladosporioides* MD2, the genes *dbat, bapt* and *dbtnbt*, which encodes 3′-N-debenzoyltaxol N-benzoyltransferase, were not detected in the transcriptome data [41]. It is unclear from Miao and colleagues whether the original sequence for the *dbat* gene published for *C. cladosporioides* MD2 in 2009 by their laboratory, is present in the genome, other than to say that no transcript was found [41]. Moreover, another *dbat* sequence for the basidiomycete-fungus, *Grammothele lineata*, published by Das, Rahman, et al., 2017 [42], does not seem to be present in the draft genome of the fungus, published later that year [43]. While in some reports the *dbat* gene has been detected, their sequences were thought to be non-specific amplification and BLASTn searches produced no significant hits, or simply the *dbat* sequences were not published in GeneBank [36-38]. Thus, our negative results obtained during the initial PCR screening using the published primers for *dbat* and *bapt* would be consistent with those obtained for published fungal genomes, that is, genes with significant homology to some of the plant genes involved in Taxol biosynthesis, are absent [40, 44].

In addition to *Annulohypoxylon* sp., other fungal genera identified included *Alternaria* sp., *Aspergillus* sp., *Fusarium* sp., *Mucor* sp., *Penicillium* sp., and *Trichoderma* sp., which is consistent with the published literature on the diversity of endophytic fungi isolated from *Taxus wallichiana* [7, 8]. Given that metabolite production can vary according to culture conditions, these fungi were not discarded yet and will continue to be evaluated for the production of bioactive compounds.

Though further analysis is required to elucidate the molecular structure of the compounds present in isolate *MUS1* showing antibacterial and antioxidant properties, our results underscore the importance of preserving the Himalayan yew in Nepal, as they harbor microorganisms that could be the source of new bioactive compounds in the future.

## Conclusion

Among the 16 endophytic fungal isolates, the *Annulohypoxylon* sp. strain *MUS1* not only showed the ability to produce Taxol, but also the production of antimicrobial and antioxidant compounds. The crude extract of the isolate *MUS1*, significantly inhibited the fluconazole-resistant yeast, *Candida albicans*, and also to a range of Gram-positive and Gram-negative bacterial pathogens, and as well has radical scavenging activity as shown by the assays. Once these bioactive compounds have been identified, they could be further developed to be used by the pharmaceutical and food industries. Thus, our results underline the importance of preserving *Taxus* spp., seeing that their fungal endophytes could be the source for the new antimicrobial and antioxidant compounds of the future.

## List of abbreviations

ATCC: (American Type Culture Collection)
AWD: (Agar Well Diffusion)
BLAST: (Basic Local Alignment Search Tool)
BLASTn: (Nucleotide BLAST)
ITS: (Internal Transcribed Spacer)
IUCN: (International Union for Conservation of Nature)
MEGAX: (Molecular Evolutionary Genetics Analysis Version X)
MHA: (Muller Hinton Agar)
MHB: (Muller Hinton Broth)
MUS1: (Endophytic fungal isolate MUS1)
NCBI: (National Center for Biotechnology Information)
PCR: (Polymerase Chain Reaction)
PDA: (Potato Dextrose Agar)
PDB: (Potato Dextrose Broth)
UV-Vis: (Ultraviolet-Visible)

## DECLARATIONS

### Ethics approval and consent to participate

Not applicable.

### Consent for publication

Not applicable.

### Availability of data and materials

The sequence obtained in this study was deposited in the NCBI GenBank database (Accession number: MN699475).

### Author contributions

DPG designed the study, carried out the majority of research activities, wrote the Manuscript. HV and AA equally contributed to conducting research activities and manuscript editing. JO, KL, ME are involved in molecular analysis of isolated and purified endophytes and editing the manuscript. MRGG participated in designing the research, guiding DPG to carry out the research and provided financial support. All authors read and approved the final manuscript.

## Acknowledgments

We would like to acknowledge the Department of Biotechnology, Kathmandu University (KU), Nepal for the facilitation of the research works. We would also like to acknowledge Mr. Basant Awasthi, Teaching Assistant, Department of Geomatics Engineering, KU for his contribution to making the location map of Mustang. Further, we want to convey warm regards to Dr. Janardan Lamichhane, Department of Biotechnology, KU and Dr. Rajani Shakya, Department of Pharmacy, KU for the HPLC analysis.

