## Supplemental figures and table for "*Annulohypoxylon* sp. strain *MUS1*, an Endophyte isolated from *Taxus wallichiana* Zucc. produces Taxol and Other Bioactive Metabolites"

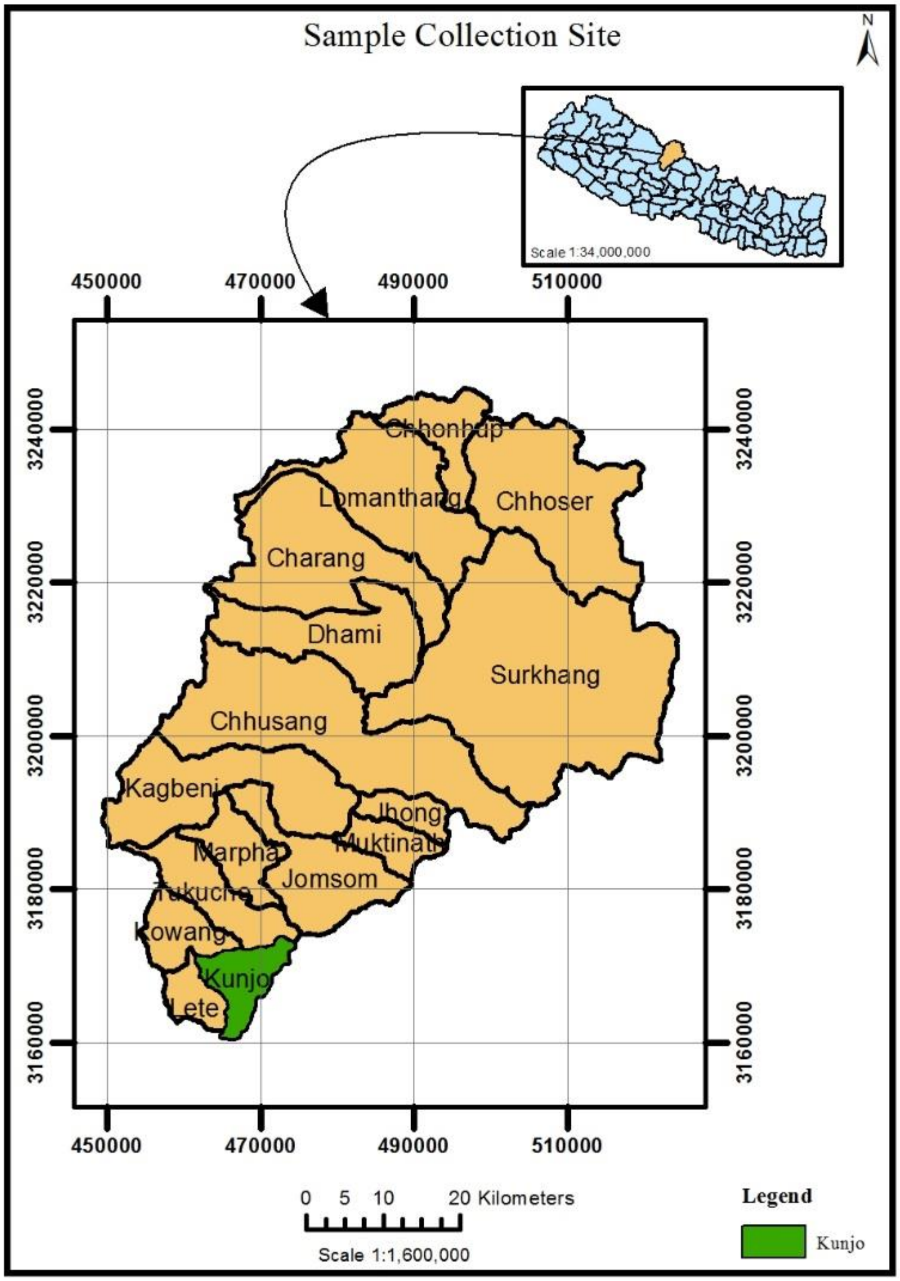


Figure S1: Location map for the sample collection site. *Taxus* plant materials were collected from *Taglung* forest area of the Lower-Mustang region in *Kunjo* VDC in the Mustang district of Nepal.

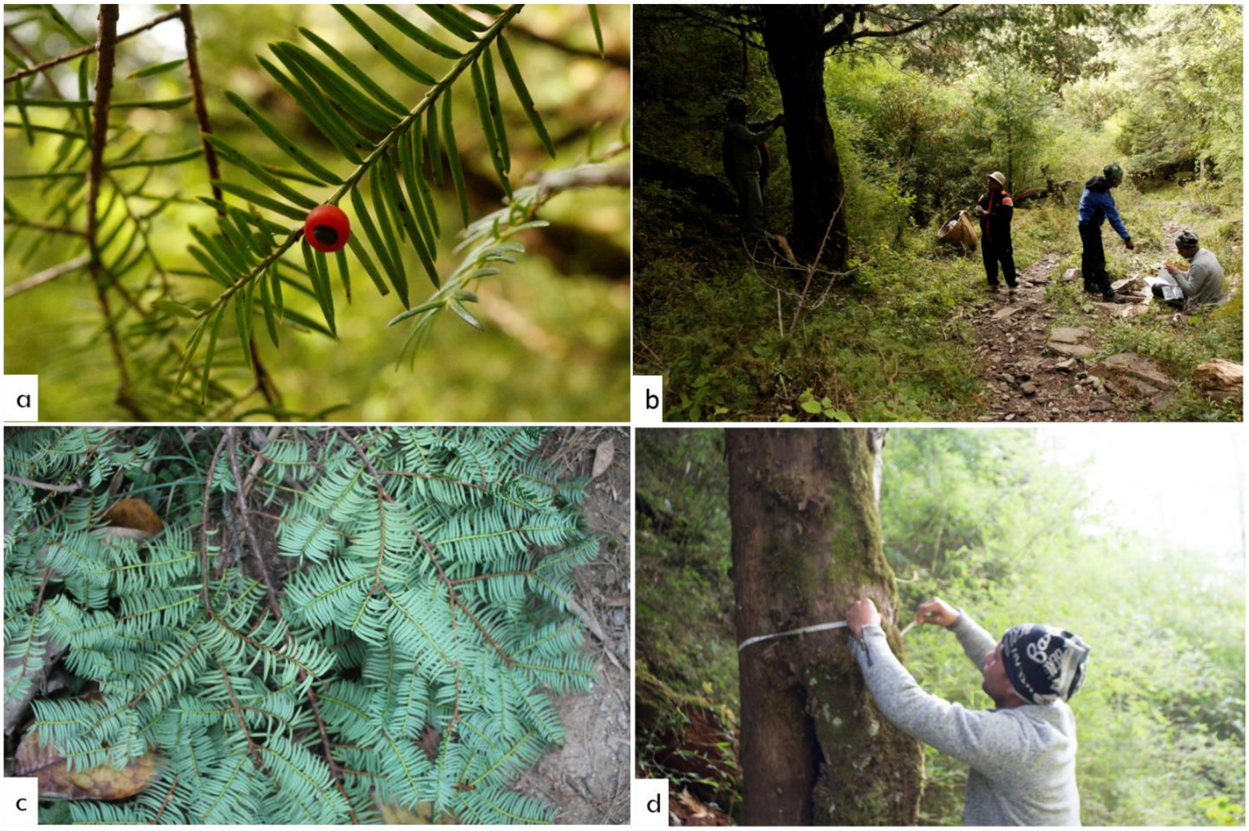


Figure S2: Sampling activities of a research group in *Taglung* forest area of the Lower-Mustang region in *Kunjo* VDC in the Mustang district of Nepal. (a) *Taxus* Aril, (b) DPG giving instruction for sample collection, (c) Twigs and Needles of *Taxusi*, and (d) AA in sample collection process.

Table S1: Sequence submitted to NCBI for *Annulohypoxylon* sp. strain *MUS1* with accession no. MN699475.

| |
|---|
| TACCGAAACGGTCTCCGTTTGGTGACCAGCGGAGGGATCATTACTGAGTTATCAA<br>AAACTCCAACCCTTTGTGAACCTACCTATGTTTCCTCCGGCGTACCGCTGTAGCCT<br>ACCCGCAGGGCTCCCCCTTAGGGGGGTTTTGCTGGGGAGGTGCCTGAGTGCTACC<br>TATCCTTCGGGGTACGGTTAGTGCAGTGAAGGTGCCGACCAAGGCCTCGGCGGCG<br>CCGAGTAGGACCGCTCCAAACTTAAGCACTTAGTGCATCCAACCCCGCGTTGAAC<br>AACTATCGAAAATTTGCTTTTGCTTTTTTTCTTTACGCTAAAACGTCTTTCCCCGGT<br>TGGAATTATTGCTCGAAATGATAATTTCCTTACCCTGTAGTCGTTTGTTTTCAAGCT<br>ACAATATATCTGCTCGAATAAAATTGCTTCAATATTTGCTCGAAAATTGTTCAAAG<br>CTCTGAGGGGTCTGAATGAATTCATAAAATTGGCAAAAGCCACCTATAAACTACG<br>GTTTTTAGGGGGTGATCAAACCAAGGTTTTAAAAACCAAATACGTTAAAACTTTC<br>AACAACGGATCTCTTGGTTCTGGCATCGATGAAGAACGCAGCGAAATGCGATAAG<br>TAATGTGAATTGCAGAATTCAGTGAATCATCGAATCTTTGAACGCACATTGCGCCC<br>ATTAGTATTCTAGTGTGCATGCCTATTCGAGCGTCATTATAACCCTTAAGCCTTGTT<br>GCTTAGCGTTGGGAATCTACCCCTCACTGAGGGGTAGTTCCTTAAATGTAGTGGCG<br>GGGTTATAGCACACTCTAAGCGTAGTAGTTTAACTCGCTTTCAGGGAGGCTGTAGC<br>TGCTTGCCGTAAAACCCCTTATAACTTATAGTGGTGACCTCGGATATGGTAGTAAT<br>GCTCCTGCTCAC |
